# Efficient genome-wide sequencing and low coverage pedigree analysis from non-invasively collected samples

**DOI:** 10.1101/029520

**Authors:** Noah Snyder-Mackler, William H. Majoros, Michael L. Yuan, Amanda O. Shaver, Jacob B. Gordon, Gisela H. Kopp, Stephen A. Schlebusch, Jeffrey D. Wall, Susan C. Alberts, Sayan Mukherjee, Xiang Zhou, Jenny Tung

## Abstract

Research on the genetics of natural populations was revolutionized in the 1990’s by methods for genotyping non-invasively collected samples. However, these methods have remained largely unchanged for the past 20 years and lag far behind the genomics era. To close this gap, here we report an optimized laboratory protocol for genome-wide capture of endogenous DNA from non-invasively collected samples, coupled with a novel computational approach to reconstruct pedigree links from the resulting low-coverage data. We validated both methods using fecal samples from 62 wild baboons, including 48 from an independently constructed extended pedigree. We enriched fecal-derived DNA samples up to 40-fold for endogenous baboon DNA, and reconstructed near-perfect pedigree relationships even with extremely low-coverage sequencing. We anticipate that these methods will be broadly applicable to the many research systems for which only non-invasive samples are available. The lab protocol and software (*“WHODAD”*) are freely available at www.tung-lab.org/protocols and www.xzlab.org/software, respectively.

---

The capacity to generate genetic data from low-quality or non-invasively collected samples, first developed in the 1990’s^1,2^, revolutionized the study of genetics, evolution, behavior, and ecology in natural populations. These methodological advances facilitated phylogenetic and phylogeographic analyses of difficult-to-sample taxa^3-5^; helped define the role of admixture in mammalian evolution^6–8^; and enabled theoretical expectations about paternal investment, kin recognition, and reproductive skew to be empirically tested, sometimes for the first time^9–12^. They also yielded important insights into the genetic viability and future prospects of threatened or endangered populations from which invasive samples are impossible to obtain^13-17^. Non-invasive genetic analysis has thus changed the ways we study population, ecological, and conservation genetics. Indeed, these fields would look very different today—and we would know far less about many species—without it.

However, techniques for non-invasive genetic analysis have changed little in the past twenty years. Collection of genetic data from non-invasively collected tissues (e.g., feces, hair, urine) continues to be labor-intensive, time-intensive, and vulnerable to technical artifacts such as allelic dropout and cross-contamination^18,19^. Further, current methods ultimately yield very small amounts of data by today’s standards. Typical studies genotype only a dozen to several dozen microsatellite loci per individual – a trivial amount compared to the data sets now routinely generated using standard high-throughput sequencing approaches. Thus, while existing methods are sufficient for basic pedigree construction and estimating some population genetic parameters (although usually with substantial uncertainty), they are severely underpowered for many other types of analyses, such as identifying signatures of natural selection, reconstructing population history and demography, and testing for genetic associations with phenotypic variation^20–22^. Similarly, analyses that require local (i.e., gene- or region-specific) information on genetic diversity, structure, or ancestry instead of genome-wide averages cannot be conducted^23-27^. Finally, because non-invasively collected genotype data are most often based on microsatellites, they cannot take advantage of new tools designed specifically for single nucleotide variants^28–30^.

Generating genome-scale data sets from non-invasive samples is challenging for two reasons. First, in many cases, the DNA extracted from these samples is low quality and highly fragmented. Second, it contains large proportions of non-host DNA. For example, only about 1% of DNA extracted from fecal-derived samples is endogenous to the donor animal (most is microbial)^31^. Sequence capture methods, in which synthesized baits are used to enrich for pre-specified target sequences from a larger DNA pool^32^, present a potential solution to both of these problems. Because shearing is a required step in library preparation, the problem of working with highly fragmented samples is obviated. Indeed, Perry and colleagues^31^ were able to target and sequence 1.5 megabases of the chimpanzee genome from fecal-derived DNA, using a modified version of sequence capture, with very low genotyping error rates relative to blood-derived DNA. More recently, Carpenter et al.^33^ reported a method for performing genome-wide sequence capture from low-quality ancient DNA samples, which recapitulate many of the challenges posed by non-invasive samples (e.g., highly-fragmented DNA and low proportions of endogenous DNA).

However, while considerable investment in single samples often makes sense in ancient DNA studies, the low levels of post-capture enrichment associated with currently available protocols are not cost-effective for population studies of non-invasive samples. Substantially higher rates of enrichment, particularly in non-repetitive regions of the genome, will be essential to overcome this limitation. In addition, computational methods for analyzing the resulting data are also required, especially given that genome-scale sequencing efforts for such samples are likely to produce low coverage data. For example, current paternity assignment approaches^34–36^ were not designed to deal with uncertain genotypes, an inevitable component of analyzing low coverage sequencing data. Thus, for capture-based methods to become broadly accessible, the development of appropriate new computational approaches is also essential.

Here, we report an optimized laboratory protocol for genome-wide capture of endogenous DNA from non-invasively collected samples, combined with a novel computational approach to reconstruct pedigree links from the resulting data (implemented in the program *WHODAD*). We validate both our lab methods and computational tools using non-invasively collected samples from 54 members of an intensively studied wild baboon population in the Amboseli basin of Kenya^37^. We also demonstrate the generalizability of our methods to non-invasive samples collected using different methods from a different baboon species from West Africa. Our protocol is cost effective, has manageable sample input requirements, yields good capture efficiency for high complexity, non-repetitive elements, and minimizes the need for extensive PCR amplification. Importantly, we find that genotype data generated from fecal samples closely match data from high quality blood-derived DNA samples from the same individuals, and provide near-perfect information on pedigree relationships even with extremely low per-sample sequencing coverage (mean = 0.49x genome coverage). Together, these methods will enable population, conservation, and ecological genetic analyses of natural populations to again take a major leap forward, into the genomic era. At the same time, they will also introduce valuable new systems to the genomics community.

## RESULTS

### DSN digestion during bait construction increases library complexity

Our protocol relies on *in vitro* transcription of biotinylated RNA baits to capture host-specific DNA from the mixed pool of host, environmental, and microbial DNA extracted from non-invasive samples. Similar to Carpenter et al^33^, RNA baits are generated from DNA templates obtained from a high quality DNA sample (here, DNA extracted from blood). This approach avoids the high cost of custom bait synthesis (as in Gnirke et al.^32^ and Perry et al.^31^), but can also produce a bait set that includes a large proportion of low complexity, repetitive regions. Consequently, reads generated from captured DNA cannot be uniquely mapped, lowering the protocol’s efficiency relative to using a more diverse bait set. To address this concern, we incorporated a novel duplex specific nuclease (DSN) digestion in the bait construction step (Fig. S1A; see Methods). Sequencing the DNA bait templates prior to *in vitro* amplification demonstrates that including the digestion step reduces the percentage of baits synthesized from low complexity/highly duplicated regions. Specifically, a 4 hour incubation of sheared DNA at 68°C followed by a 20 minute DSN digestion in the presence of human Cot-1 greatly improved the efficiency of capture, producing the highest complexity bait library of the five conditions we tested. Compared to DNA templates from a non-DSN-digested library, bait templates produced using these conditions reduced the number of reads mapping to multiple locations by 2.6-fold (from 19.2% to 7.5%; Fig. S2).

### Capture-based enrichment

We validated our full capture protocol (bait construction followed by capture of endogenous DNA and sequencing of captured fragments) using fecal-derived DNA (fDNA) samples collected from 54 individually recognized yellow baboons (36 males and 18 females; Fig. 1) from the Amboseli baboon population, an intensively studied population in which maternal and paternal pedigree relationships are known for a large set of individuals^9,37,38^. We produced data for 52 of the samples in two successive capture efforts: “Capture 1” was conducted on fDNA from 24 baboons, and “Capture 2” was conducted on fDNA from 28 additional baboons after making multiple improvements to our initial protocol (changes to the protocol between capture efforts are described in detail in Table S1; Table S2 provides detailed information on sequencing coverage and mapping statistics). Data from the remaining two individuals, “LIT” and “HAP”, were generated to compare the captured fDNA sample with data derived from sequencing high-quality genomic DNA samples (gDNA) extracted from blood for the same individuals.

**Figure 1.**
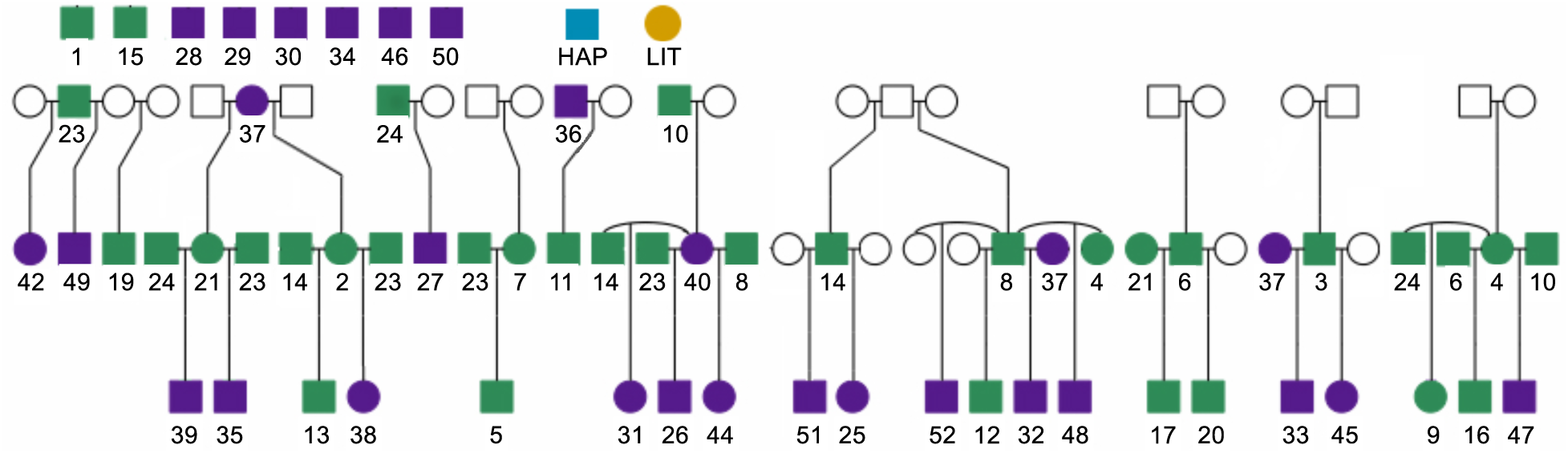
Pedigree of a subset of baboons monitored by the Amboseli Baboon Research Project. Samples from both males (squares) and females (circles) were enriched in Capture 1 (green) or Capture 2 (purple). Unfilled circles/squares represent baboons that connect individuals in our pedigree, but who were not sequenced as part of this study. Each sequenced individual is represented by a unique number (below the circle/squares); note that some individuals are repeated in the figure because baboons often produce offspring with multiple mates. The paired fDNA and gDNA samples came from two individuals, HAP (blue) and LIT (orange), who were members of the study population but are not connected to this pedigree.

Our protocol (Fig. S1) resulted in substantial enrichment of baboon DNA in the post-capture versus pre-capture samples (see Table S2 for sample-specific details). A mean of 44.56% (range: 10.28-83.17%) of post-capture fragments mapped to the baboon genome, despite starting with pre-capture samples that contained a mean of only 2.04% endogenous baboon DNA, as estimated by qPCR (range 0.19-8.37%). However, in Capture 1 a large proportion of the mapped fragments were identified as PCR duplicates (mean_capture1_=71.97% of mapped fragments, range_capture1_: 51.43-88.46%; Fig. 2A). After removing PCR duplicates, a mean of 9.16% of the post-capture reads in Capture 1 were non-duplicate mappable fragments (range_capture1_=2.23%-23.75%), producing a mean coverage of 0.20x per sample relative to the mappable baboon genome (mean sequencing depth of 5.8 Gb per sample; range_capture1_ = 0.04-0.49x; Fig. 2B). These numbers translated to an overall mean fold enrichment of 39.8-fold for mapped reads (range_capture1_: 8.0-111.8-fold, s.d.=25.2), and 9.6x enrichment of non-PCR duplicate mapped reads (range_capture1_:3.9-22.4-fold, s.d.=5.0; Fig. 2C).

**Figure 2.**
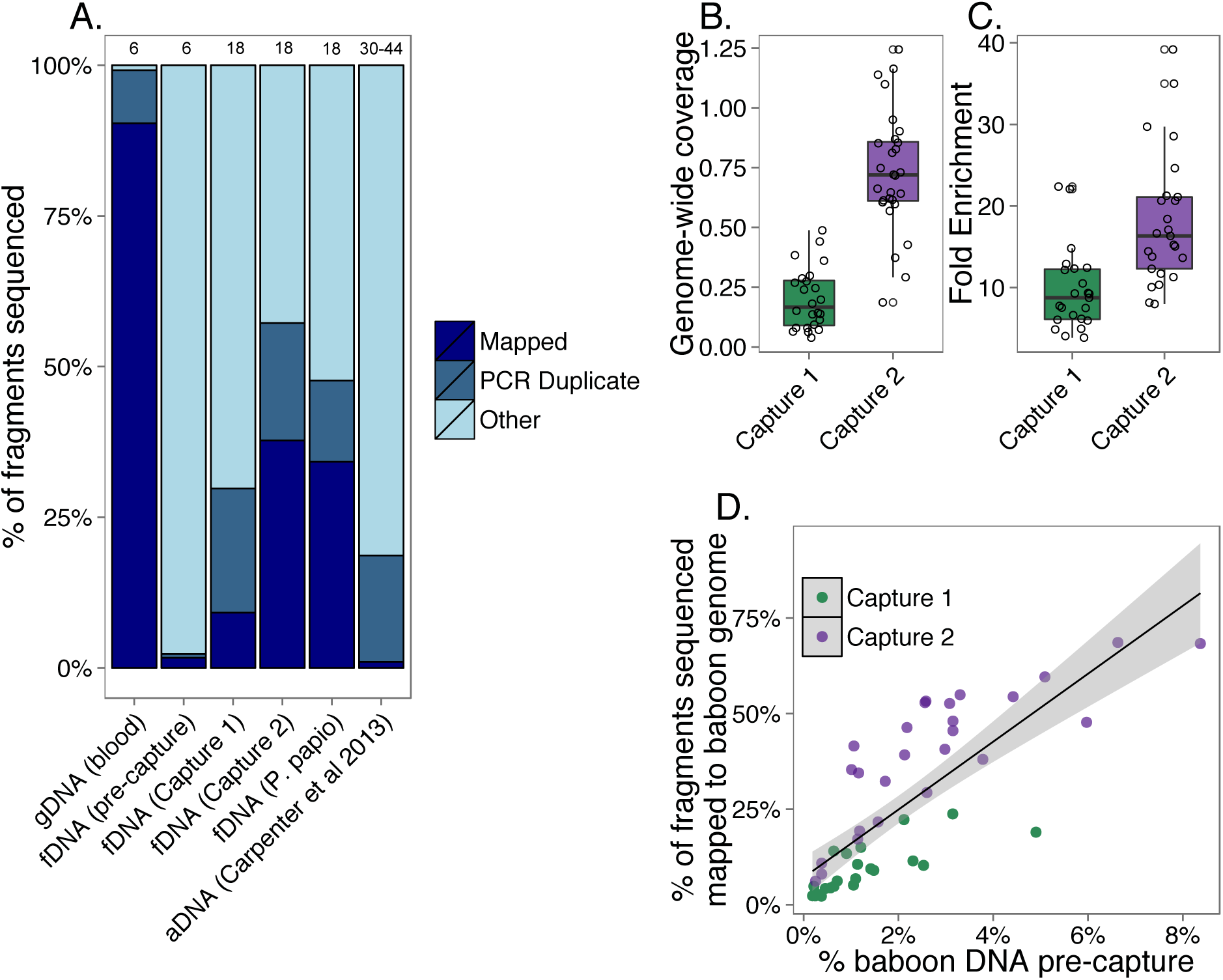
fDNA enrichment results. (A) Percent of sequencing reads that mapped to the baboon genome and were not PCR duplicates (“Mapped:” dark blue); mapped and were PCR duplicates (“PCR Duplicate:” blue); or did not map and likely represent environmental or bacterial DNA in the case of fDNA/aDNA and unmappable fragments in the case of gDNA (“Other:” light blue). “gDNA” represents genomic DNA derived from the blood samples for LIT and HAP; “aDNA” represents ancient DNA data from capture-based enrichment reported in Carpenter et al^33^. Numbers above each bar show the total number of PCR cycles used in each protocol. (B) Capture 2 produced significantly greater genome coverage than Capture 1, despite similar number of overall reads generated per sample (two-sample t-test, T=9.7, p=3.0×10^−12^). On average in Capture 2, we obtained ~0.73x coverage of the genome with 5.76Gb of sequencing. If all 5.76Gb mapped to the baboon genome as non-PCR duplicates, we would have produced ~2.2x genome-wide coverage. (C) Capture 2 also produced significantly greater fold enrichment of baboon DNA (fold enrichment is measured as % non-duplicate baboon DNA post-capture divided by % baboon DNA pre-capture: two-sample t-test, T=4.4, p=7.3×10^−5^). (D) The amount of baboon DNA in the sample pre-capture (% baboon DNA pre-capture, based on qPCR of the single copy *c-myc* gene^39^) is strongly correlated with the percentage of baboon fragments obtained in post-enrichment sequencing (Pearson’s r=0.80, p=1.0×10^−11^). However, even samples with low amounts of endogenous DNA (<2%) exhibit substantial fold enrichment using our protocol (mean_capture1_=10.60x, mean_capture2_=24.82x).

Based on our results for Capture 1, we made multiple protocol improvements prior to conducting Capture 2 (Table S1). The improved protocol was twice as effective on average, resulting in a mean 18-fold enrichment of high quality, analysis-ready reads and a maximum fold enrichment of close to 40-fold (range_capture2_ = 8.0-39.2-fold; Fig. 2C; by comparison, methods optimized for ancient DNA achieved a mean of 5.5-fold enrichment of non-PCR duplicate fragments^33^; Fig. 2A). Specifically, the protocol changes improved the proportion of non-duplicate mapped fragments by more than four-fold, from a mean proportion of 9.16% in Capture 1 to a mean proportion of 37.74% in Capture 2 (range_capture2_=6.16-68.61%) and reduced the proportion of PCR duplicates among mapped reads two-fold (from 71.97% in Capture 1 to 36.97% in Capture 2). This improvement translated to an increase in overall genomic coverage from a mean of 0.20x in Capture 1 to 0.73x in Capture 2 (mean total sequencing of 5.7Gb per sample; range_capture2_ = 0.19-1.24x; Fig. 2B). This improvement in coverage was not explained by increased sequencing depth in Capture 2 (Table S2). Thus, while we would need to sequence a pre-capture fDNA sample 50-100 times as deeply as a blood or tissue-derived sample to produce the same level of coverage, our capture method reduces this difference to approximately 2 times the sequencing effort. Importantly, our method was also successful in enriching fDNA samples (n=8) from independent samples collected from Guinea baboons (*P. papio;* Fig. 2A, Table S2), suggesting that our results are highly generalizable across different species and storage and extraction methods.

### Sample attributes influencing capture efficiency

The amount of baboon DNA in the pre-capture fDNA sample was the strongest predictor of enrichment success. Specifically, the percent of baboon DNA pre-capture, as assessed via qPCR, was positively correlated with the percentage of non-duplicate fragments mapped post-capture (Fig. 2D; T=6.88, p=1.72×10^−8^). Samples from Capture 2 had more pre-capture baboon DNA than samples used in Capture 1 because we attempted to optimize the input samples based on our initial analyses in Capture 1 (Capture 1 mean = 1.21%, range = 0.19-4.90%; Capture 2 mean = 2.80%, range = 0.25-8.37%). However, even when controlling for this difference, enrichment of samples from Capture 2 was improved over Capture 1. This pattern is observable whether assessed using the percent of baboon DNA fragments sequenced post-capture (T_capture2_=10.00, p=6.76×10^−13^) or fold enrichment relative to pre-capture amounts (T_capture2_=6.89, p=1.69×10^−8^), and could not be explained by differences in the length of sequence fragments or overall sequencing depth (Fig. S3; Table S2). The amount of fDNA library used in the capture reaction was also weakly positively correlated with the percent of baboon DNA fragments sequenced post-capture, after controlling for the amount of baboon DNA in the pre-capture sample (T_ng_fDNA_library_=2.09, p=0.042; Table S2).

### Library complexity, distribution of reads, and GC content

The post-capture libraries included a higher proportion of PCR duplicates relative to reads generated from high-quality genomic DNA samples, for which fewer rounds of PCR amplification were required (PCR duplicate proportion: mean_fDNA_capture1_=69.6%, mean_fDNA_capture2_=36.8%, mean_gDNA_=11.3% of mapped reads; 18 rounds of PCR in the capture protocol versus 6 for the high-quality samples). For comparison, this proportion is much lower than reported for aDNA samples, which go through more rounds of PCR amplification (mean_aDNA_=94.6%; Fig. 2A and Fig. S4^33^). Despite increases in clonality, the number of non-duplicate reads continued to increase with increasing sequencing depth, with the slope of this relationship especially favorable for Capture 2 (Fig. 3). Thus, deeper sequencing of post-capture libraries should continue to increase genome-wide coverage, albeit not as efficiently as sequencing blood-derived gDNA samples.

**Figure 3.**
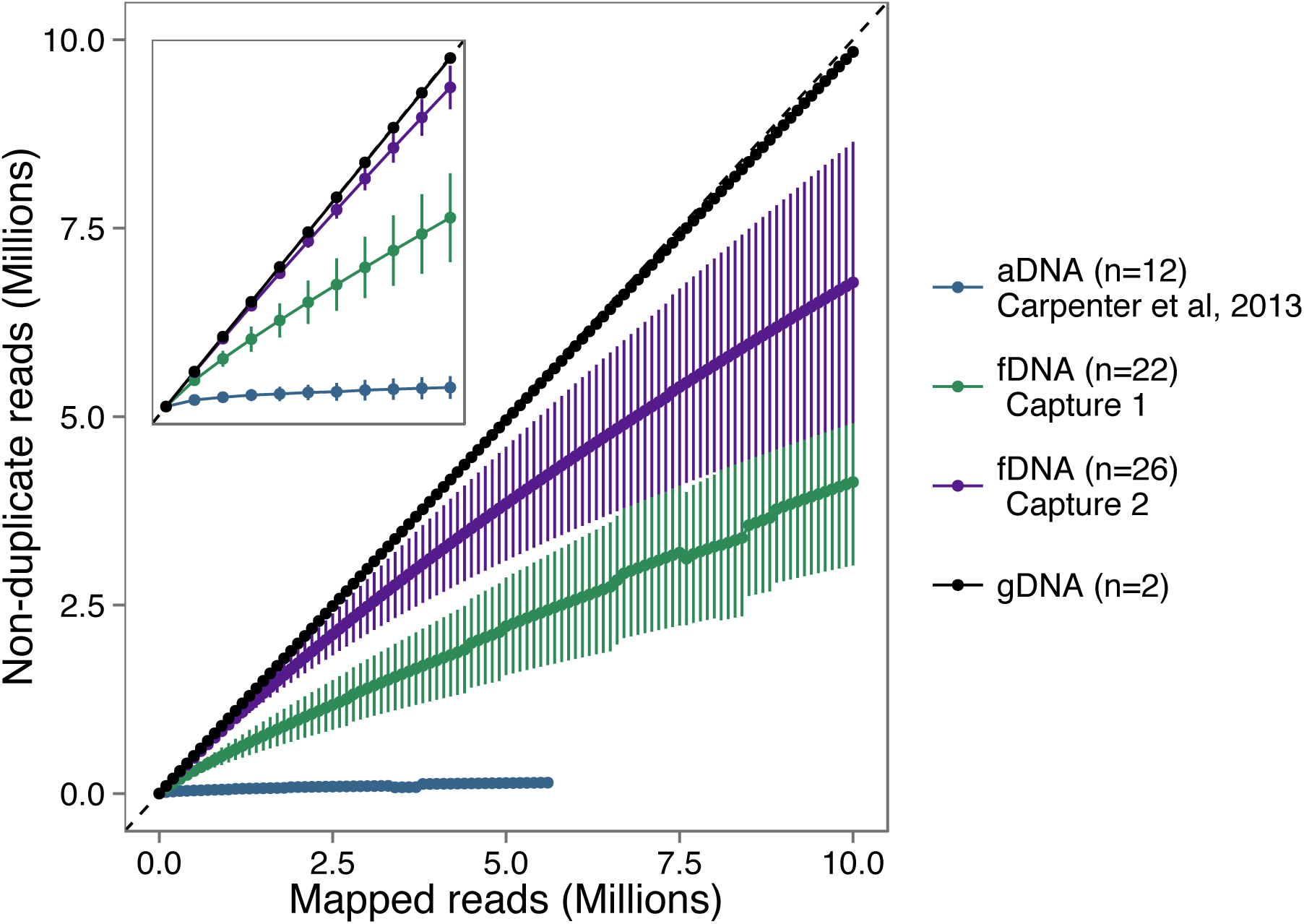
Increased sequencing effort produces increased numbers of non-duplicate reads. The number of mapped reads plotted against the number of non-duplicate reads mapped (mean ± SD; plotted using the program “preseq”^40^). More complex libraries (i.e., those containing more non-duplicate fragments) have a slope closer to 1 (as in the case of the gDNA libraries), while less complex libraries have a shallower slope and asymptote at a smaller value. The main plot shows the first ten million mapped reads for each sample. The inset shows the same plot for the first million mapped reads.

As with other capture-based methods^33,41^, a modest fraction of the mapped fragments mapped to the mitochondrial genome (mtDNA). When we included all mapped reads, this fraction was similar in libraries from Capture 1 and Capture 2 (mean_capture1_=6.55%; mean_capture2_=6.73%; Fig. S5A). However, Capture 2 resulted in significantly more unambiguously non-duplicate mtDNA-mapped reads than Capture 1, largely due to the paired-end sequencing used in Capture 2 (mean_capture1_=0.47% of all mapped reads; mean_capture2_=6.46%; Fig. S5B). The higher number of non-duplicate mtDNA reads in Capture 2 thus produced much deeper overall coverage of the mitochondrial genome (Fig. S5C), despite the fact that the ratio of mtDNA to nuclear DNA mapped reads was comparable between the two captures (Fig. S5D). Finally, the distributions of read GC content for post-capture reads using our protocol, the DNA template for the RNA baits, and aDNA libraries were highly similar (Fig. S6). This observation suggests that any GC bias relative to the genome appears during bait construction and/or sequencing, not during the hybridization step.

### Post-capture fDNA-derived genotype data are consistent with individual identity and independently established pedigree relationships

To assess the accuracy of genotypes called from post-capture fDNA libraries, we compared genotype data from paired blood-derived gDNA (without capture) and post-capture fDNA libraries for two individuals, LIT and HAP. Using genotypes for sites that were called with a genotype quality (GQ) > 20 in both the fDNA and gDNA data sets for either LIT or HAP, we found that the majority of the genotypes called in both data sets were concordant (86.5%, or 270,724 of 312,739 sites for the LIT paired samples; 77%, or 30,948 of 40,132 sites for the HAP paired samples; note that we had lower coverage for the HAP fecal-derived sample than for the LIT fecal-derived sample). As expected, the majority of the discordant sites occurred when the low coverage fDNA sample was called as homozygous and the high coverage gDNA sample was called as heterozygous (77.7% of discordant LIT_gDNA_ heterozygous sites; 74.4% of discordant HAP_gDNA_ heterozygous sites). Importantly, the fDNA genotype captured at least one of the alleles from the gDNA genotype in 99.8% (LIT) and 99.6% (HAP) of these discordant sites. Thus, even when genotypes called in fDNA and gDNA samples from the same individual were discordant, they were almost always compatible.

Further, we found that genotypes called from the post-capture fDNA libraries were more similar to the genotypes called from their high-quality gDNA counterparts than they were to other post-capture fDNA libraries. Specifically, *k0* values from *lcMLkin*^42^, which estimate the probability that two samples share no alleles that are identical by descent, were much smaller for the LIT_fDNA_-LIT_gDNA_ paired samples (0.487) and HAP_fDNA_-HAP_gDNA_ paired samples (0.243) than for *k0* values calculated for the two blood-derived samples when compared to any other fDNA sample (*k0* range LIT_fdna_ versus other fDNA samples = 0.996 − 1.000; Z=849.2, p < 10^−20^; *k0* range HAP_fdna_ versus other fDNA samples = 0.786 − 0.999; Z=10.6, p < 10^−20^; Fig. 4A).

**Figure 4.**
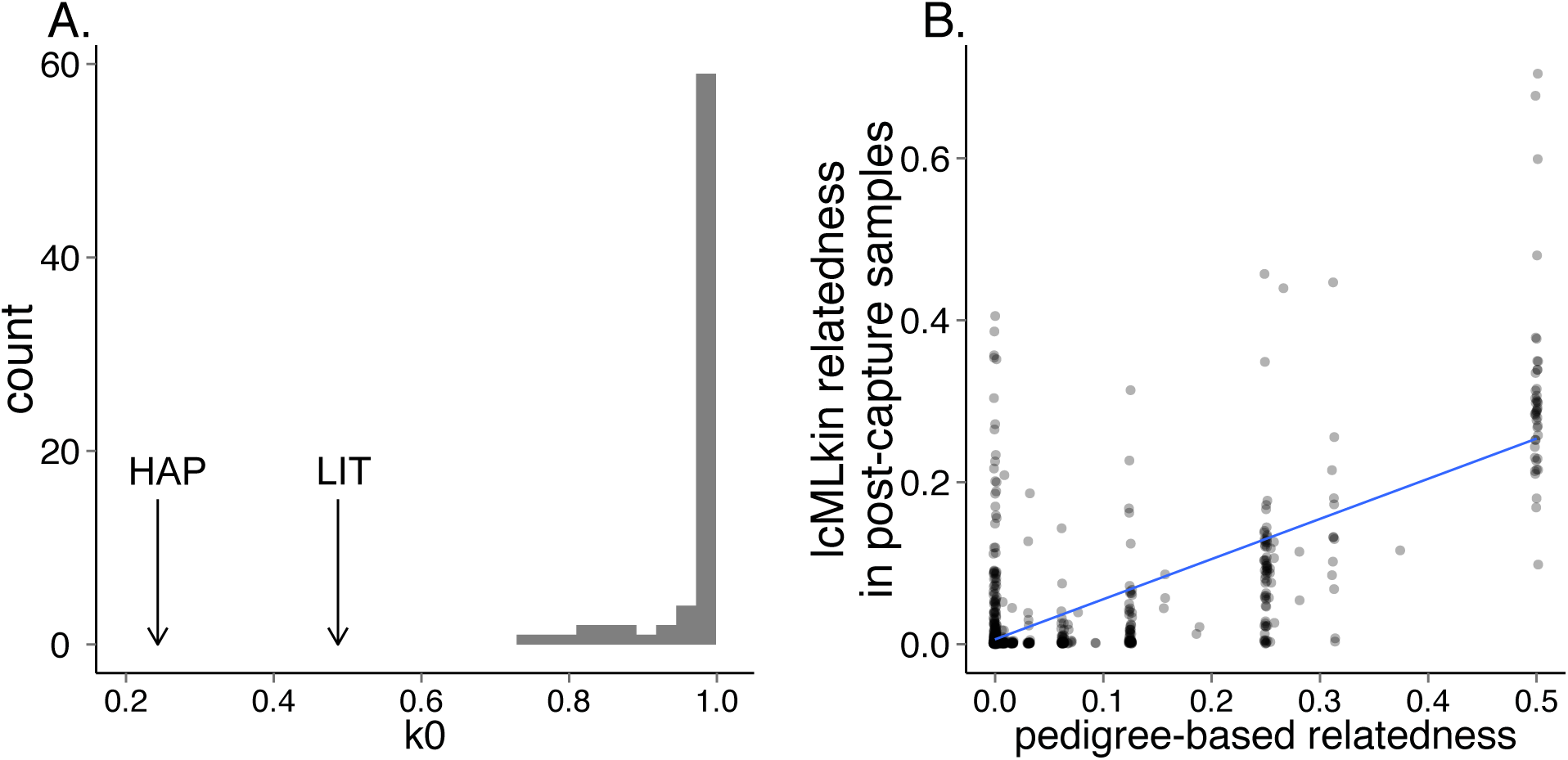
Post-capture genotype data are consistent with individual identity and pedigree relationships. (A) The *k0* values for the HAP and LIT fDNA-gDNA paired samples (arrows) were significantly lower than the range of *k0* values for LITfDNA and HAP_fDNA_ versus any other fDNA sample (gray distribution). Lower *k0* values reflect increased relatedness (i.e., decreased probability of no IBD sharing). (B) Estimated dyadic relatedness values were correlated with independently obtained pedigree relatedness values calculated using the R package *kinship2* (Sinnwell et al. 2014; r=0.73, p<10^−16^). Both *k0* and the estimated relatedness values were calculated with *lcMLkin*^42^.

For the 48 extended pedigree individuals (Fig. 1, including 8 Amboseli baboons with no known relatives in the pedigree), we then tested if estimated relatedness values from *lcMLkin*^42^ in the post-capture data were correlated with relatedness values obtained from the independently constructed pedigree (based on known mother-offspring relationships and microsatellite-based paternity assignments: see Methods). Using a filtered set of 127,654 single nucleotide variants (see Methods for filtering parameters), we found a strong correlation between the two measures (Pearson’s r=0.73, p<10^−16^; Fig. 4B). This correlation improved further if we imposed thresholds for the minimum number of sites genotyped in both individuals (“shared sites”) in a dyad (Fig. S7). For example, if we removed all dyads with fewer than 2,000 shared sites (84 of 1,128 dyads, or 7.4%), the correlation between pedigree relatedness and genotype similarity reached r=0.86 (p<10^−16^).

### Paternity inference using WHODAD

Current methods for assigning paternity (e.g., CERVUS^34,35^ and exclusion^36^) assume genotype certainty, such that individuals are assigned a deterministic genotype at each locus (i.e., 0, 1, or 2, or a microsatellite repeat number; while a low level of measurement error, i.e., due to lab handling, can be modeled, this error rate is held constant across genotype calls). This assumption is violated in low coverage sequencing data, in which genotypes are not known with certainty and this uncertainty varies across genotype calls. However, the relative *probabilities* of each genotype can be estimated, given estimated population allele frequencies and sequencing coverage information. To conduct paternity inference and pedigree reconstruction in this context, we therefore developed a novel approach to integrate information across low coverage sites, implemented in the program *WHODAD.* Our method has two components. The first component identifies a top candidate male and tests whether he is significantly more related to the offspring than any other candidate male, using a p-value criterion. The second component tests whether the dyadic similarity between the top candidate and offspring is consistent with a parent-offspring dyad, using posterior probabilities obtained from a mixture model (see Methods and Fig. S8).

Using *WHODAD,* we assigned paternity to all father-offspring pairs (n = 27) represented in the independently established extended pedigree in Fig. 1. Note that our approach represents a particularly conservative test because it departs from the usual practice of first identifying a likely set of candidate fathers based on demographic and prior pedigree information (the approach used in producing the pedigree in Fig. 1). For 15 of the 27 offspring, we produced genotype data from the known mother with our enrichment protocol. *WHODAD* identified the same father as shown in the pedigree in 12 of these 15 trios (80%); in the other 3 trios (20%), no candidate male satisfied *WHODAD*’s paternity assignment criteria (in all three of these cases, sequencing coverage was very low for either the pedigree-identified father or offspring: 0.04-0.17x). For the remaining 12 offspring, we did not generate genotype data using our enrichment protocol for their mothers (i.e., their mothers were not among the samples run in Capture 1 or 2). To test all 27 father-offspring dyads together, we therefore re-ran *WHODAD* excluding maternal genotype information. In this setting, *WHODAD*’s paternity assignments agreed with the pedigree data in 22 of 27 (81%) cases (Fig. 5). Notably, when the pedigree-identified father was included in the data set, *WHODAD* never assigned paternity to a different male, whether or not maternal genotype data were available. Because our method is highly robust to exclusion of maternal genotype data, we therefore performed all subsequent analyses assuming maternal genotype data were *not* available. This approach allowed us to evaluate all father-offspring dyads, and also captures a scenario that may often occur in studies of natural populations.

**Figure 5.**
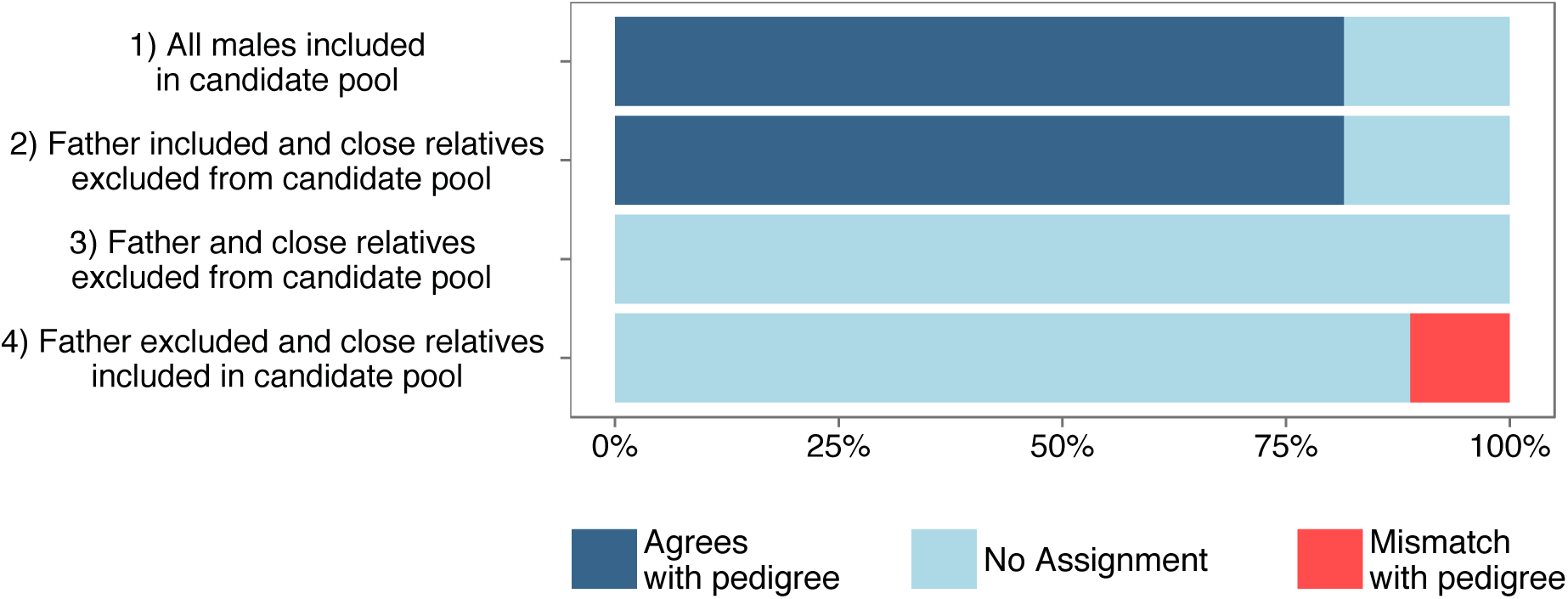
Paternity inference with *WHODAD* using low coverage genotype calls. 1) When all males (n=34) were included in the pool of candidate fathers (top bar), *WHODAD* assigned paternity to the same father identified in the pedigree for 22 of 27 (81%) of offspring (see assignment criterion in Methods; dark blue). The remaining offspring were not assigned a father based on *WHODAD*’s assignment criteria (5 of 27; light blue). 2) *WHODAD*’s accuracy was identical when we removed all close male relatives of the offspring (r ≥ 0.25) from the pool of candidate fathers. 3) When we removed all close relatives, including fathers, from the candidate pool, no fathers were assigned, as expected. 4) Finally, when we removed the father from the candidate pool but retained close relatives, our method incorrectly assigned paternity to 11% of offspring (3 of 27; bottom bar). All three incorrectly assigned fathers were closely related to the offspring (in two cases the assigned father was the half-brother of the offspring and in one case the assigned father was the son of the offspring).

The presence of close relatives, such as full or half-siblings, can influence the accuracy of paternity assignment if these close relatives are also included as candidate fathers^35,44-46^. Thus, to examine how the presence of close male kin influenced the accuracy and confidence of *WHODAD*’s paternity assignments, we conducted three additional analyses. First, when all close male kin were removed from the candidate list of potential fathers (r ≥ 0.25), but the father was retained, our method performed equivalently to the case when both father and close relatives were in the candidate pool. Second, when we removed all close male kin *including* the father, none of the best candidate fathers from the conditional probability analysis (0%) were assigned as fathers based on *WHODAD*’s assignment criteria (Fig. 5). Third, when we removed the father from the pool of candidate fathers, but included close male kin, 11 % of the best remaining candidates (3 of 27 cases) were incorrectly assigned as fathers, based on comparison to the pedigree (Fig. 5). All 3 of these false positives were close male relatives: in two cases *WHODAD* assigned the half-brother of the offspring as the likely father, and in one case *WHODAD* assigned the son of the offspring as the likely father. The best balance between maximizing the number of true positives while minimizing the number of false positives was achieved by combining both the p-value and mixture model criteria (see Methods). This approach outperformed either component used alone (Fig. S9). For example, when all males were included in the candidate pool, the combined approach resulted in an 81% true positive rate and a 0% false positive rate, while just using the *k0* values in a mixture model resulted in the same true positive rate (81%), but an additional 11% false positive rate (Fig. S9).

## DISCUSSION

Our capture-based method strongly enriches the proportion of host DNA in low-quality DNA extracted from feces (fDNA). Our method is the first use of genome-wide enrichment-based capture methods^33,47,48^ for non-invasively collected samples, which represent a major resource for behavioral, conservation, and evolutionary genetic studies in natural populations. Importantly, our protocol increases efficiency and lowers cost by reducing the input requirements and number of PCR cycles relative to previous methods^31^ and, in our final protocol, achieves up to 40-fold enrichment of post-capture endogenous DNA relative to pre-capture levels. We also show, for the first time since Perry et al^31^, that capture libraries from low-quality samples produce genotype data that are highly concordant with genotype data derived from high-quality, non-captured samples from the same individuals.

We anticipate that data generated through this protocol could be leveraged for a wide variety of applications. To illustrate this point for paternity analysis, one of the most central components of genetic studies in natural populations, we present an accompanying method, *WHODAD,* that produces results in near-perfect concordance with an independently constructed pedigree, using low-coverage data generated with our enrichment protocol (note that the few cases in which assignments could not be confidently made could readily be addressed with slightly deeper sequencing coverage, similar to typing more markers in conventional microsatellite analysis). By incorporating demographic and behavioral data often used to constrain pedigree reconstruction, as well as prior information about other pedigree links, its performance would be improved even further. For instance, in reconstructing pedigree links in the Amboseli population, we generally include only plausible candidates (e.g., we exclude males who were immature or not yet born at the offspring’s conception), not all males with genotype data, as we did here.

Together, these results provide valuable, accessible wet lab and computational tools for moving studies of difficult-to-sample natural populations forward into the genomics era. Importantly, our methods can be generalized to produce low complexity DNA-depleted RNA baits for any species in which at least one high-quality DNA sample is available (or potentially a closely related species^48^).

### Costs of performing the protocol

At the time of publication, using the same reagents as we used here and sourced from the same locations, the cost of generating these data is ~$60/sample (including sequencing costs). Importantly, our method does not require the commercial synthesis of targeted capture probes, which is a relatively expensive step for many capture-based approaches^31,32^. Thus, the majority of the costs are accounted for by the streptavidin-coated Dynalbeads ($11/prep), RNA baits ($5/sample) and High Sensitivity Bioanalyzer chips for quality control ($9/sample). Replacing Ampure XP beads with homemade SPRI beads would reduce the per-sample costs considerably, as would pooling adapter-ligated fDNA samples prior to hybridization (instead of post-hybridization, as reported here). For a multiplexed pool of 10 samples, we estimate that using these two strategies would result in a per-sample cost of ~$29. Indeed, we have verified that multiplexing samples prior to hybridization does not result in loss of capture efficiency, and actually resulted in improved yield of mapped, non-PCR duplicate reads (~61% of reads; mean of 117-fold enrichment, range = 54.8 − 257.2-fold; Fig. S10A), although it did result in more uneven coverage of samples sequenced within a pool (Fig. S10B). Multiplexing also has the advantage of reducing the amounts of input DNA per sample and the number of PCR cycles required for the initial library preparation step. We are currently pursuing improvements to the protocol along these lines.

Based on achieving 40% non-PCR duplicate, mapped reads after capture (the mean result for Capture 2 samples), we estimate that the sequencing costs of a 1x genome for baboon (~2.9 Gb) would be about $200 (based on paired-end, 125-bp sequencing at $2,000 per lane and exclusion of PCR duplicates). This cost per sample is approximately twice the cost of genotyping 14 microsatellites from the same fDNA sample—the previous strategy for the main study population, the Amboseli baboons^49^— but provides substantially more genetic information. These estimates will drop further as the cost of high-throughput sequencing continues to fall, making application of our approach to whole populations increasingly feasible. Notably, our finding that useful sequencing reads do not asymptote with deeper sequencing (Fig. 3) also suggests the feasibility of producing a high-quality, high-coverage genome from such samples if one were to sequence more deeply than required for the analyses reported here.

Finally, to make the current protocol as cost-effective as possible, we recommend that researchers use qPCR quantification to choose DNA samples with the highest proportion of host DNA possible—the strongest predictor of the foldchange enrichment in endogenous DNA post-versus pre-capture (Fig. 2D).

### Assigning paternity using WHODAD

The lack of available tools for working with low coverage genomic data— realistically, one of the most likely data types to be produced for studies of natural populations—represents a major barrier to moving from low-throughput marker genotyping to genome-scale analyses. The pedigree structure of a study population is fundamental to understanding its genetic structure and social organization. However, current methods for pedigree reconstruction are unable to cope with high levels of genotype uncertainty. The approach we have implemented in WHODAD takes this uncertainty into account, suggesting one simple application for the wet lab methods presented here. Indeed, our method performed well when compared to an independently constructed extended pedigree, with its major challenges—differentiating between close relatives in a candidate pool—comparable to those reported for existing software^34,35,45,46^. Importantly, while analyses of pedigree structure using previously available methods are greatly aided by prior knowledge of mother-offspring relationships^34^, maternal links do not appear to be necessary for WHODAD analyses, which performs well even when no maternal information is available (Fig. 5; Fig. S8).

### Conclusions

High-throughput sequencing approaches solve one problem of working with low-quality, non-invasive samples: the sheared nature of the original samples. Capture approaches have demonstrated great promise for solving the second major problem— large proportions of non-endogenous DNA—since the results published by Perry et al (2010). Motivated by parallel work on ancient DNA, our results help to fulfill this promise by providing methods to perform cost-effective scaling of sequence capture from non-invasive samples on a genome-wide scale, coupled with analytical methods to deal with the resulting data. Our protocols add an important tool to the range of available options for genetic data generation from such samples. Notably, for questions in which investigators are specifically interested in variants in *a priori-defined* subsets of the genome (e.g., the exome^50,51^), targeted capture with synthesized baits, followed by much deeper sequencing, may still be the best option. However, for the many types of analyses that use genome-scale data (e.g., local ancestry analysis; genome-wide scans for selection, including in non-coding regions; reconstruction of population demographic history^20-27,30^), our approach will be more useful, especially as the costs of high-throughput sequencing continue to fall.

Here, we focused specifically on DNA obtained from fecal samples, which are one of the most commonly collected types of non-invasive samples: they contain information not only about host genetics, but also about endocrinological parameters^52^, gut microbiota^53^, parasite burdens^54^, and, as recently demonstrated for human infants, gene expression levels^55^. The sample banks already available for many natural populations thus open the door to population and evolutionary genomic studies in species in which such analyses were previously impossible. As the costs of data generation continue to fall, and the limiting factor for many studies becomes high quality phenotypic data, we envision that such studies will rapidly move far beyond the simple analyses of paternity and pedigree structure reported here.

## Methods

### Bait generation

Similar to Carpenter et al^33^, we use a cost-effective *in vitro* synthesis method based on T7 RNA polymerase amplification of sheared DNA from a high-quality sample (Fig. S1A). We extracted genomic DNA from a blood sample collected from an olive baboon (*Papio anubis*) who was unrelated to any of the individuals in the samples we wished to enrich. To generate baits, we sheared 5 μg of purified DNA to a mean fragment size of 150 bp, and then end repaired and A-tailed the fragments using the KAPA DNA Library Preparation Kit for Illumina Sequencing. We purified the resulting reaction using a 1.8x ratio of AMPure beads to sample volume.

We annealed custom adapters to the A-tailed library by incubating the following reagents for 15 minutes at 20 °C: 10 μL 5x ligation buffer (KAPA Biosystems); 5 μL DNA Ligase (KAPA Biosystems); 1 μL 25 μΜ custom adapter; ≤34 μL of A-tailed DNA; and H_2_O up to 50μl total volume. The custom adapters we used (EcoOT7dTV: Fwd 5’-GGAAGGAAGGAAGAGATAATACGACTCACTATAGGGCCTGGT, EcoOT7dTV: Rev 5’-/5Phos/CCAGGCCCTATAGTGAGTCGTATTATCTCTTCCTTCCTTCC) differ from those used in other protocols^33,47,48^. Specifically, they contained: 1) a T7 RNA polymerase recognition site; 2) flanking sequence that improves T7 transcription efficiency^56^; and 3) an Eco0109I restriction enzyme cut site that allowed us to later cleave off the adapter sequence from T7 amplified RNAs (rather than blocking it, as in Carpenter et al.^33^).

We then digested the purified, adapter-ligated DNA with duplex-specific nuclease (DSN; Axxora). DSN is a Kamchatka crab-derived enzyme that specifically degrades double-stranded DNA but not single-stranded DNA, allowing us to take advantage of DNA reassociation kinetics to reduce the representation of repetitive regions in the bait set (Fig. S2)^57^. We performed DSN digestion in fifteen 2 μL aliquots, each mixed with 1 μL 4x hybridization buffer (200 mM HEPES pH 7.5; 2 M NaCl; 0.8 mM EDTA) and 1 μL human Cot-1 DNA (1 μg/μL). We denatured the DNA by heating to 98°C for 3 minutes, held the reaction at 68°C for 4 hours, and then added 4 μl H_2_O, 1 mL 10x DSN Buffer, and 1 μl DSN (1 U/μL) to the reaction. After 20 minutes of digestion, we stopped the reaction by adding 5 μL 2x DSN Stop Solution (10 mM EDTA) and purified it with 2.4x AMPure beads.

Next, we used Klenow DNA polymerase to blunt end the non-digested DNA, size-selected for 200 - 300 bp fragments on a 2% agarose gel, and purified the size-selected fraction using the Zymoclean Gel DNA Recovery Kit (Zymo Research). After purification the aliquots were PCR amplified for 16 cycles using 25 μL 2x HiFi Hot Start ReadyMix (KAPA Biosystems) and 1 μL each of 25 μΜ primers EcoOT7_PCR1 (5’-GGAAGGAAGGAAGAGATAATACGACTCACT) and EcoOT7_PCR2 (5’-TACGACTCACTATAGGGCCTGGT). Following amplification the bait DNA libraries were purified using 1.8x AMPure beads and the resulting product was visualized on a Bioanalyzer DNA 1000 chip (Agilent Technologies).

Finally, we *in vitro* transcribed the DNA libraries to construct biotin-tagged RNA baits using the MEGA Shortscript Kit (Life Technologies) and Biotin-UTP (Illumina). Briefly, 125-150 nM of DNA baits were incubated at 37°C for 4 hours in the following reaction: 2 μL T7 10x reaction buffer, 2 μL each of T7 ATP, GTP, CTP, and UTP solutions (75 mM), 1 μL Biotin-UTP (50 mM), 2 μL T7 enzyme mix, and water to 20 μL total volume. We then digested the DNA template by adding 1 μL TURBO DNase (Life Technologies) to the reaction and incubating at 37°C for 15 minutes. We purified the resulting reaction with the MEGAClear Transcription Clean-Up Kit (Life Technologies) and eluted in a final volume of 70 μL. To cleave off the adapter sequence, we digested the RNA baits with the *Eco*O1091 enzyme (NEB). Lastly, the baits were again purified with the MEGAclear Clean-Up Kit, eluted in 70 μL, and quantified on a Bioanalyzer RNA 6000, Eukaryote Total RNA chip (Agilent Technologies).

### Samples, DNA extraction, and qPCR quantification

Baboon samples were stored in 95% ethanol and fDNA was extracted using the QIAamp DNA Stool Mini Kit (Qiagen; with slight modifications as described in Alberts et al.^38^), or using the QIAxtractor (protocol available here: http://amboselibaboons.nd.edu/assets/84050/alberts_fecal_genotyping_protocol_sca.docx)^38^. The majority of the sampled individuals (48 of 54) were either members of a single extended pedigree or were unrelated males living in the same study population that were genotyped for inclusion in pedigree building/paternity testing for members of that pedigree (Fig. 1). For LIT and HAP, gDNA was extracted from blood samples using the Qiagen Maxi Kit (Qiagen).

To assess our protocol’s generalizability to samples collected and stored using different methods, we also extracted fDNA samples from 8 unhabituated Guinea baboons *(P. papio)* sampled in West Africa. These samples were stored in either 90% ethanol or soaked in 90% ethanol and then dried using silica beads (i.e., the “two-step” method^58,59^). They were then extracted using either the Qiagen DNA Stool Mini Kit or the Gen-ial First DNA All Tissue Kit (Table S2).

We assessed the proportion of endogenous DNA in each fDNA sample using qPCR against the *c-myc* gene, as described in Morin et al.^39^.

### Library preparation

All samples were fragmented to the desired size (200 or 400 base pairs: see Table S1) using a Bioruptor instrument (Diagenode). Illumina sequencing libraries were then generated from the fragmented DNA using either the KAPA DNA library kits for Illumina (Capture 1) or NEBNext DNA Ultra library kit (Capture 2: see Table S1). Libraries were amplified for 6 PCR cycles prior to capture-based enrichment. Sample-specific details of library preparation and sequencing results are described in Table S1. Note that we changed several steps between Capture 1 and Capture 2 based on interim improvements in the protocol (also detailed in Table S1). Because the methods used in Capture 2 were ultimately more effective, the updated Capture 2 protocol is described in the Methods except where explicitly noted.

### Capture-based enrichment

We modified the capture methods from Gnirke et al^32^ and Perry et al^31^ (Fig. S1B). For each capture, we hybridized 121 – 626 ng of the fDNA libraries generated as described above to the RNA baits. First, we mixed each fDNA library with 2.5 μL human Cot-1 DNA (1 mg/mL), 2.5 μL salmon sperm DNA (1 mg/ml), and 0.6 μL index-blocking reagent (“IBR”, 50 μM). This mixture was incubated for 5 minutes at 95°C followed by 12 minutes at 65°C. Next, we added 13 μL of hybridization buffer (10x SSPE, 10x Denhardt’s solution, 10 mM EDTA, 0.2% SDS, preheated to 65°C), 7 μL hybridization bait mixture (1 μL SUPERase-In, 750 ng RNA baits, and water up to a total volume of 7 μL, preheated to 65°C) to the fDNA mixture, and incubated the complete mixture at 65°C for 48 hours (see Fig. S11 for comparison of alternative bait concentrations and incubation times).

After incubation, we purified the enriched fDNA sample using 50 μL Dynal MyOne Streptavidin T1 beads (Invitrogen). To do so, the beads were washed a total of three times with 200 μL binding buffer (1 M NaCl, 10 mM Tris-HCl [pH 7.5], 1 mM EDTA) and resuspended in 200 μL of binding buffer. Next, the entire fDNA/RNA hybridization mix was added to the 200 μL Dynal MyOne Streptavidin T1 bead and binding buffer slurry. We incubated this mixture at room temperature for 30 minutes on an Eppendorf Thermomixer at 700 rpm. The mixture was placed on a magnetic rack, the supernatant was discarded, and the beads were washed once with 500 μL low stringency wash buffer (1x SSC, 0.1% SDS) followed by a 15-minute incubation at room temperature. The beads were then washed three times with 500 μL high stringency wash buffer (0.1x SSC, 0.1% SDS) with a 10 minute room temperature incubation between each wash. After the final wash, the enriched fDNA fraction was eluted from the beads with 50 μL elution buffer (0.1 M NaOH), transferred to a new tube containing 70 μL “neutralization buffer” (1 M Tris-HCl, pH 7.5), purified with 1.8x AMPure beads, and eluted in a 30 μL volume. A final PCR was carried out in a 50 μL reaction volume using 23 μL of the post-hybridization fDNA and either: 1) 25 μL 2x KAPA High Fidelity master mix and 2 μL TruSeq universal primer (Capture 1); or 2) 25 μL 2x NEBNext High Fidelity PCR master mix, 1 μL universal PCR primer, and 1 μL NEB indexing primer (Capture 2). After 12 PCR cycles the final reaction was purified with 1x AMPure beads, eluted in 20 μL H_2_O, and visualized on a Bioanalyzer High Sensitivity DNA chip.

### Sequencing and alignment

All high-throughput sequence generation was conducted on the Illumina HiSeq platform (see Table S1 for sequencing details). The resulting sequencing reads were mapped to a *de novo* assembly of the *Papio cynocephalus* genome (alignment available at https://abrp-genomics.biology.duke.edu/index.php?title=Other-downloads/Pcyn1.0) using the default settings of the *bwa mem* alignment algorithm v0.7.4-r385^60^. Duplicate reads were marked and discarded in subsequent analyses using the “MarkDuplicates” function in PicardTools (http://picard.sourceforge.net). To facilitate comparison across samples of differing coverage, and because coverage of the gDNA samples was much higher (~30X) than for the fDNA samples for LIT and HAP (1.4 and 0.27 respectively), we downsampled the gDNA libraries to 0.73x coverage (the median coverage of samples in Capture 2) using “DownsampleSam” in PicardTools.

### Comparison sequencing data sets

In several analyses, we compared our capture-based enrichment results to two independent datasets: i) a previously published capture-based enrichment of ancient DNA (aDNA) samples (NCBI SRA accession: SRP042225)^33^, and ii) shotgun sequencing from six Capture 1 fDNA samples prior to hybridization (“pre-capture”; Table 1). The aDNA samples were aligned to the human genome *(hg38)* and the pre-capture fDNA samples were mapped to the *de novo P. cynocephalus* genome assembly.

### Library complexity, distribution of reads, and GC content

We calculated the complexity of each library using two methods. First, we used the ENCODE Project’s PCR Bottleneck Coefficient (PBC), which calculates the percent of non-duplicate mapped reads out of the total number of mapped reads^61,62^. The PBC ranges from 0 to 1, where more complex libraries have higher numbers. Second, we used the function “c_curve” from the program *preseq* (v1.0.2) to plot the number of non-duplicate fragments mapped vs. the number of total mapped fragments^40^. More complex libraries (i.e., those with fewer duplicate fragments) have a c_curve slope closer to 1, meaning that increasing sequencing depth continues to provide novel information. Less complex libraries have a shallower slope and asymptote at smaller values. Lastly, we evaluated the GC bias for each sequencing library using Picard Tools, “CollectGCBiasMetrics” (http://picard.sourceforge.net).

### Sample attributes influencing capture efficiency

To determine the sample attributes that predicted the success of our capture protocol, we first modeled the relationship between the proportion of non-duplicate reads that mapped to the baboon genome after capture (our primary measure of protocol success) and (i) the percent of endogenous baboon DNA in the pre-capture samples; (ii) the amount of fDNA library (ng) that went into the capture; and (iii) whether the sample was captured using our initial protocol or the second version of the protocol (i.e., in “Capture 1” or “Capture 2”). Second, we investigated the relationship between the same three variables and a secondary measure of protocol success, the fold-change enrichment of baboon DNA in the sample pre-versus post-capture. Pre-capture concentrations of endogenous DNA in fDNA samples were measured as the concentration of baboon DNA estimated using qPCR, relative to the concentration of total DNA estimated using the Qubit High Sensitivity fluorometer (Life Technologies). To ensure that our qPCR-based measures were well calibrated, we confirmed the relationship between qPCR-based estimates and pre-capture sequence-based estimates of endogenous DNA in 6 samples for which both values were available (R^2^=92; Fig. S12). All statistical analyses were carried out in R^63^.

### Variant calling

We used two different approaches to call variants and genotypes in our sample: SAMTOOLS^64,65^ and the Genome Analysis Toolkit (GATK)^66-68^. In downstream analyses, we only retained variants that were identified by both methods, a strategy that produces a higher ratio of true positives to false positives than variants identified by a single method alone^69^. Duplicate-marked alignments were used as input for both methods. SAMTOOLS variant calling was carried out using *mpileup* and *bcftools,* with a maximum allowed read depth (-D) of 100. GATK variant calling was carried out following the GATK Best Practices for GATK v3.0, for variant calling from DNA-seq. To minimize potential batch effects introduced by the two capture efforts, we used the following strategy. First, we called genotypes using reads from each capture independently. Second, we re-called genotypes using reads from both captures together. Third, we extracted the union set of variants called in steps 1 and 2 for downstream analysis.

Because no reference set of genetic variants is currently publicly available for baboons, we used a bootstrapping procedure for base quality score recalibration. Briefly, we performed an initial round of variant calling on read alignments without quality score recalibration. From this variant call set, we extracted a set of high confidence variants that passed the following hard filters: quality score ≥100; QD < 2.0; MQ < 35.0; FS > 60.0; HaplotypeScore >13.0; MQRankSum < -12.5; and ReadPosRankSum < -8.0 (as described in Tung, Zhou, et al.^70^). We then recalibrated the base quality scores for each alignment using this high-confidence set as the database of “known variants” and repeated the same variant calling and filtering procedure for 3 additional rounds. Finally, we identified the intersection set between the variants called from GATK and SAMTOOLS, respectively, using the *bcftools* function isec^64^. To produce our final call set, we removed all sites that were genotyped in only one of the capture efforts, had a minor allele frequency of <0.05, or were within 10 kb of one another, using *vcftools*^71^.

### Estimating relatedness

To produce an estimate of relatedness between samples in our pedigree and to test for concordance between fecal and blood-derived samples for the same individuals, we used the program *lcMLkin*^42^. *lcMLkin* uses the genotype likelihoods generated by GATK for each genotype call to calculate two measures: (i) *k*0, the probability that two individuals share no alleles that are identical by descent, and (ii) *r*, the coefficient of relatedness^42^. Several other methods have been developed,^72,73^ to estimate relatedness from thousands of SNPs, but *lcMLkin* yielded the best match to pedigree-based estimates in our data set (Fig. S13).

We also compared genotype calls for the matched fecal and blood-derived samples using GATK’s GenotypeConcordance function^68^. This tool allowed us to determine concordance rates between data sets for different classes of variants (e.g., 0, 1, or 2). For the majority of variant sites, we expected that the genotypes would be completely concordant (i.e., the same genotype called in the fDNA and gDNA samples from the same individual). However, for calls reported as discordant, we expected that most errors would reflect cases in which the low coverage sample was called as homozygous and the high coverage sample was called as heterozygous, as low read depth makes observation of both alleles at a heterozygous site less likely.

### WHODAD: Paternity inference and pedigree reconstruction

Our paternity prediction model is based on a naïve Bayes classifier that takes advantage of the rules of Mendelian segregation within pedigrees. Using data from all sites genotyped in a potential father-mother-offspring trio or, when the mother is not genotyped, all sites genotyped in a potential father-offspring dyad, it estimates the posterior probability that a potential candidate is the true father of a given offspring.

Our approach can be broken into three steps (Fig. S8). First, we estimate, for each candidate male, the conditional probability that he is the true father of a given offspring, given the genotype data for the candidate, offspring, and mother, if known (below we show the case in which genotype information is available for the mother, but the model is similar when maternal genotype information is missing). Second, we assign a p-value for the top candidate male from the first step, for the null hypothesis that he is *not* more related to the focal offspring than the other candidates tested. Third, we calculate the probability that the genotype data for the top candidate and offspring are consistent with a true parent-offspring relationship, using a mixture model. Steps (ii) and (iii) perform subtly different functions in our analysis: (ii) tests that the top candidate is significantly more related to the offspring than any other candidate, whereas (iii) tests that the dyadic similarity between the candidate and the offspring look as expected for parent-offspring dyads. We have found that combining both approaches is key to detecting true positive fathers while minimizing false positive calls that can occur when true fathers are not in the pool of genotyped candidates (Fig. S8).

*Step 1: estimating conditional probabilities for each trio.* For a given offspring or mother-offspring dyad, our goal is to infer the true genetic father from a pool of *n* candidates. For the *i*^th^ candidate, we use data for the *L_i_* variants for which we have genotype information for the known mother-offspring dyad and for the candidate father. Assuming the true father is present in the candidate pool (i.e., he has been genotyped), the probability that the *i*^th^ potential candidate is the father is:

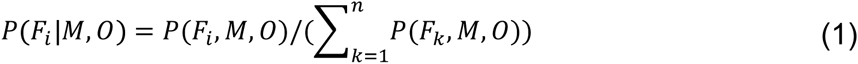
where *P*(*F_i_*|*M*, *0*) denotes the probability that the candidate is the father, conditional on the (known) mother-offspring dyad; *P*(*F_i_*, *M*, *0*) denotes the joint probability of the whole trio; and 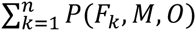 is the sum of the joint probabilities of all possible trios evaluated in the analysis. In practice, we normalize these conditional probabilities to take into account differences in the number of variants evaluated for each trio by taking the *L_i_*^th^ root:

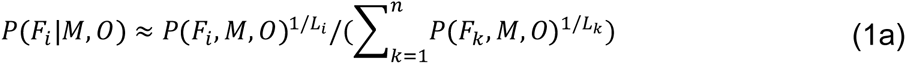
Each joint probability can be calculated in turn as:

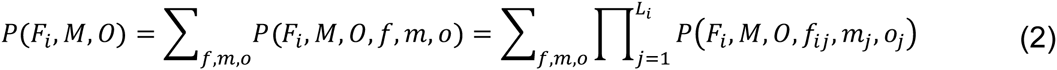
where *m_j_, f_ij_* and *o_j_* represent the genotype data for the j^th^ variant of the mother, the candidate father, and the offspring, respectively. Genotypes take values in {0, 1, 2} (i.e., the number of copies of the reference allele at each individual-site combination). Importantly, although equation (2) unrealistically assumes independence across loci, this assumption does not change the relative order of trio joint probabilities.

The probability *P*(*F_i_, M, O, f_ij_, m_j_, o_j_*) for each locus can be further decomposed as:

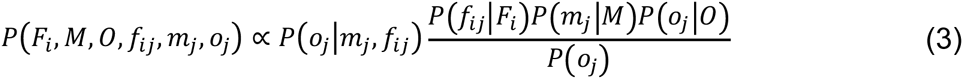
where we take genotype uncertainty into account by using GATK’s genotype probabilities to calculate the conditional genotype probabilities for *P*(*f_ij_*|*F_i_*), *P*(*m_j_*|*M*), and *P*(*o_j_*|*O*) over all possible genotype values at each site-individual combination (i.e., the probabilities that each genotype is 0, 1, or 2, which sum to 1). We also ignore the scaling constant *P*(*F_i_*)*P*(*M*)*P*(*0*) because it cancels out in the numerator and denominator of (1). The marginal probability of the offspring’s genotype, *P*(*o_j_*), is calculated from the minor allele frequency of the variant in the population. Finally, the conditional probability *P*(*o_j_*|*m_j_, f_ij_*) is based on the rules of Mendelian transmission (e.g., Marshall et al., 1998). Due to genotype uncertainty in low coverage data, the values of *P*(*F_i_*|*M, O*) are small. However, the highest value is usually assigned to the most likely father (based on comparison to the pedigree; see Results) and we can directly assess the strength of the relative evidence for the top candidate versus other candidates in Step 2 by calibrating these values against permuted data.

*Step 2: calculating resampling-based p-values.* To compute p-values for each paternity assignment, candidates are ranked based on their conditional probability *P*(*F_i_*|*M, O*) of being the true father. The log ratio of conditional probabilities between the highest probability father and the second best candidate is the test statistic:

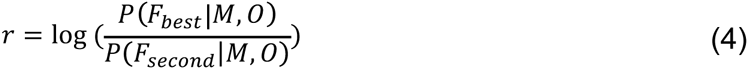

To assess significance for *r*, we then simulate genotype data for a set of *n* unrelated candidate fathers based on allele frequency information for each locus in the analysis and sequence coverage information for the real candidates, at each of the loci for which they were genotyped in the true data set. Specifically, for each locus-simulated unrelated candidate combination, *f_ij_*, where *i* indexes a (real) candidate male and *j* indexes the locus, we simulate a vector of genotype probabilities for the candidate father, (*f_ij_*_0_, *f_ij_*_1_, *f_ij_*_2_), which sum to 1. The number of probability vectors simulated for each candidate is based on the number and identity of the loci observed in the real data. For example, if the top candidate in the real data was evaluated based on 10,000 sites, we would simulate an unrelated male with genotype vector probabilities simulated for each of those 10,000 sites; if the second best candidate was evaluated at 9,000 sites, we would simulate an unrelated male with genotype vector probabilities simulated for each of those 9,000 sites; and so on. The variant sets for different simulated candidates need not be identical, and are in fact highly unlikely to be so in practice.

To simulate each vector, we draw values from a Dirichlet distribution (i.e., a distribution on probability vectors that sum to one). In principle, the Dirichlet distribution for each biallelic site could be parameterized by the genotype frequencies for each of the three potential genotype values, *Dir*(*π_j_*_0_, *π_j_*_1_, *π_j_*_2_), with genotype frequencies equal to the Hardy-Weinberg expected values based on the allele frequency of the reference allele (i.e., *p*^2^, 2p(1-p), (1-p)^2^, with p estimated from the data). However, the low coverage in our data introduces additional noise into this sampling problem, so we instead draw values from the following Dirichlet distribution:

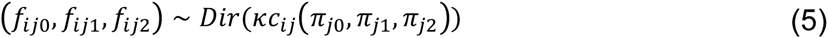
where *c_ij_* is the read depth (coverage) for the site in (true) candidate father *i,* and *κ* is a concentration parameter common to all sites and candidate fathers, estimated from the real data using the method of moments. *κ* can be thought of as a scaling factor for the effect of coverage on variance in (*f_ij_*_0_, *f_ij_*_1_, *f_ij_*_2_). To make the simulations as realistic as possible, all parameters are estimated from the real data as follows:

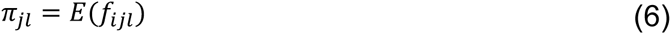
where the expectation is based on the allele frequencies for the reference allele estimated across all individuals, for each locus *j* and genotype *l* combination, and:

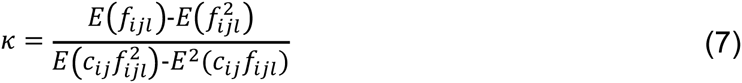
where the expectations are based on the allele frequencies (as above) across all individuals and loci, and across all 3 possible genotype values (0, 1, and 2) for each locus-individual combination. Our estimates for *π_ij_* and *κ* are based on the observed average values from the data, which approximate the expected value.

After simulating genotype data for each candidate male as if he were unrelated to the focal offspring, we can obtain a new value of *r* (equation 4) from the simulated data. By repeating this procedure s times, we can compute a p-value for the hypothesis that the best candidate in the true data is no more related to the focal offspring than any other candidate in the data set. This p-value is equal to the proportion of times the simulated test statistics exceed the observed test statistic. It intuitively corresponds to the probability of seeing a gap as large as the true gap between the conditional probabilities for the best and second best candidates, if all candidates were in fact unrelated (or equally related) to the focal offspring.

*Step 3: estimating the posterior probability of paternity. WHODAD*’s inference method, like other paternity inference methods (e.g., CERVUS^34,35^), can falsely assign paternity to a close relative if the true father is not included in the pool of potential fathers. Such false positives arise because these methods do not actually test the hypothesis that the assigned father is the true father, but rather whether the assigned father is significantly more closely related to the focal offspring than other candidates in the pool. A more direct method would be to test the probability of observing the data for a father-offspring dyad (or father-mother-offspring trio) under the *alternative* hypothesis that the assigned father is the true father. Testing the alternative hypothesis is non-trivial with low-coverage data, and by itself can also yield incorrect inferences (Fig. S9). However, in combination with the resampling-based p-values described above, it can improve paternity assignments.

To estimate the probability of the data given the best candidate-offspring dyad, we take advantage of the fact that dyadic measures of genotype similarity, relatedness, or other estimates of identity-by-descent should differ for true parent-offspring pairs compared to all other dyads (except for full sibs). By utilizing the many dyadic values in a data set of mothers, offspring, and candidate fathers, we should therefore be able to distinguish father-offspring dyads from dyads involving other relatives or unrelated pairs. Notably, this method allows us to use dyadic values for mother-offspring pairs to maximum effect.

We use a normal mixture clustering approach and the *k0* value from the R package *lcMLkin,* where low *k0* values predict high pedigree relatedness (other measures of dyadic relatedness could be substituted, but the *k0* values produced the best correlation with known pedigree-based measures of relatedness in our sample: Fig. S13). We denote y_b_ as the vector of logit-transformed *k0* measurements for the best candidate-offspring dyads for all tested father-offspring dyads; y_1_ as the vector of logit(*k0*) measurements for all known mother-offspring dyads, if any are present (y_1_ can be an empty vector if no mother-offspring dyads were sampled); and y_0_ as the vector of logit(*k0*) measurements for all other dyads. Thus, y_0_ captures the distribution of logit(*k0*) values for non-parent-offspring dyads; y_1_ captures the distribution of logit(*k0*) values for known parent-offspring dyads; and y_b_ contains a mixture of logit(*k0*) values for both true parent-offspring dyads and non-parent-offspring dyads.

We first work only with y_0_, and use a mixture model approach to assign the logit(*k0*) value for each dyad *i* into one of K component normal distributions (fit using the *mixtools* function in R, with a default value of K=5). Components with lower mean values for *k0* can be thought of as capturing the distribution of logit(*k0*) values for highly related dyads (e.g., half-siblings), whereas components with high mean values capture distantly related or unrelated dyads (if relatedness coefficients were used instead of *k0,* this direction would be reversed: low values would correspond to distantly related dyads instead). For *y*_1_, all dyads are from the same relatedness category (mother-offspring), so logit(*k0*) values in *y*_1_ can be modeled by a single distribution parameterized by a mean and a variance. Finally, for *y_b_*, values of logit(*k0*) can be assumed to be drawn from either the distribution on *y*_1_ or from one of the distributions (likely one with a low mean value) in the mixture model for *y_0_*:

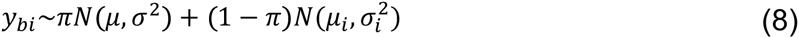
where for the *i*th individual in *y_b_*, *μ_i_* and 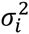 are the mean and variance for one of the distributions in the mixture model of *y*_0_; *μ* and *σ*^2^ are the mean and variance for the distribution on *y*_1_; and *π* is the probability that a value in *y_b_* belongs to the parent-offspring distribution or one of the distributions fit in the mixture model for other dyads. To infer these parameters, for each dyad in *y_b_,* we assign *μ_i_*, 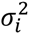 to the mean and variance of the mostly likely normal component by evaluating the likelihood under all K components. We then combine y_1_ and y_b_ to jointly infer *π, μ, σ*^2^ in equation (8).

Finally, we introduce a latent indicator variable z_bi_ for each dyad to indicate if the *i^th^* dyad in *y_b_* is a true father-offspring dyad. The probability of being a true father-offspring dyad, or P(z_bi_=1), becomes the final statistic used to assess our paternity assignments. To infer P(z_bi_=1), we use an expectation-maximization algorithm (see Supplementary Methods for detailed information about the EM steps). WHODAD considers a male as the likely true father of a focal offspring if he was (i) the candidate with the highest conditional probability of paternity; (ii) assigned a p-value from our simulations < 0.05; and (iii) P(z_bi_=1) > 0.9.

### Testing the accuracy of paternity assignment using WHODAD

We assigned paternity using the methods detailed above for all previously identified father-offspring pairs (n = 27) in the Amboseli pedigree (Fig. 1). This pedigree was constructed using a combination of observational life history data on female pregnancies and infant care (to infer maternal-offspring dyads), demographic data to identify possible candidate fathers, and microsatellite genotyping data analyzed in the program CERVUS (with confidence >95%; see Alberts et al.^38^ for additional detail).

Our data set contained maternal genotype information derived from the fecal enrichment protocol for 15 of these individuals (56%). We first used *WHODAD* to assign paternity for these 15 offspring while incorporating the genotype data from their mothers. To assess the accuracy of *WHODAD* in the absence of maternal genotype data, we then repeated the paternity analysis for the same 15 offspring without including the mother’s genotype. For this analysis, we were also able to include the 12 additional offspring for whom we did not have genotype data from the mother, but had genotype data from the known father (n=27).

To examine how the presence of close male kin influenced the accuracy and confidence of *WHODAD*’s paternity assignments, we conducted three additional analyses. First, to assess the accuracy of *WHODAD* when the pedigree-assigned father is the only close male relative present, we removed all close relatives of the offspring except the father (r≥0.25, e.g., grandfathers, half-sibling or full-sibling brothers) from the pool of potential fathers. Second, to test if *WHODAD* assigned a father with high confidence even when no close relatives were present, we removed all close male relatives, including the pedigree-assigned father, from the pool of candidate males. Third, to assess the risk of confidently (but erroneously) assigning a close male relative as the likely father when the pedigree-assigned father was not genotyped, we removed the father from the pool of potential fathers. For all *WHODAD* analyses we report assignment accuracy based on whether the father was identified by *WHODAD* with a p-value less than 0.05 and a P(z_bi_=1) > 0.90. Offspring were not assigned a father (“no assignment”) when the best candidate male was identified with a p-value > 0.05 or a P(z_bi_=1) < 0.90.

## ACKNOWLEDGEMENTS

We would like to thank the Kenya Wildlife Service, Institute of Primate Research, National Museums of Kenya, National Council for Science and Technology, members of the Amboseli-Longido pastoralist communities, Tortilis Camp, and Ker & Downey Safaris for their assistance in Kenya. We also thank Jeanne Altmann and Elizabeth Archie for their generous support and access to the Amboseli Baboon Research Project data set and samples; Raphael Mututua, Serah Sayialel, Kinyua Warutere, Mercy Akinyi, Tim Wango, and Vivian Oudu for invaluable assistance with the Amboseli baboon sample collection; Emily McLean for assistance in identifying samples from the extended pedigree; and Tauras Vilgalys for assistance in drawing the pedigree. For access to the Guinea baboon samples, we thank Julia Fischer, Dietmar Zinner and José Carlos Brito; the Wild Chimpanzee Foundation for logistical support in Guinea; and the Ministère de l’Environnemnt et de la Protection de la Nature and the Direction des Parcs Nationaux in Senegal, the Opération du Parc National de la Boucle du Baoulé and the Ministère de l’Environnement et de l’Assainissement in Mali, the Office Guinéen de la Diversité Biologique et des Aires Protégées and the Ministère de L’Environnement, des Eaux et Forêts in Guinea, and the Ministère Délégue auprès du Premier Ministre, Chargé de l’Environnement et du Développement Durable in Mauritania. Finally, we thank PJ Perry, Luis Barreiro, Greg Crawford, Tim Reddy, and members of the Alberts and Tung labs for their feedback on earlier versions of this work. This work was supported by National Science Foundation grants DEB-1405308 (to JT) and SMA-1306134 (to JT and NSM). GHK was supported by the German Academic Exchange Service (DAAD), the Christiane-Nüsslein-Volhard Foundation, The Leakey Foundation, and the German Primate Center. XZ was supported by a grant from the Foundation for the National Institutes of Health through the Accelerating Medicines Partnership BOEH15AMP.

## DATA ACCESSIBILITY

All resequencing data sets reported in this manuscript will be deposited in the NCBI Short Read Archive (SRA) upon acceptance. A reviewer URL for the metadata and run info for these data sets is available at ftp://ftp-trace.ncbi.nlm.nih.gov/sra/review/SRP064514_20151007_093529_a3f2a910685f5b07f5f45a5fc1fdb389.

## AUTHOR CONTRIBUTIONS

JT, NSM, SM, and XZ conceived and designed the research. MLY, AOS, JBG, GHK, and NSM performed all laboratory experiments. SAS, JT, and JDW provided the genome assembly. WHM, SM and XZ developed the computational methods. WHM and XZ implemented the software. NSM, JT, and XZ analyzed the data. SCA and GHK provided samples, reagents, and logistical support. NSM, JT, and XZ wrote the manuscript with input from all of the coauthors.

## COMPETING FINANCIAL INTERESTS

The authors declare no competing financial interests.

